# Scaling Dense Representations for Single Cell with Transcriptome-Scale Context

**DOI:** 10.1101/2024.11.28.625303

**Authors:** Nicholas Ho, Caleb N. Ellington, Jinyu Hou, Sohan Addagudi, Shentong Mo, Tianhua Tao, Dian Li, Yonghao Zhuang, Hongyi Wang, Xingyi Cheng, Le Song, Eric P. Xing

**Author notes:** Work done during internship at GenBio AI.

## Abstract

Developing a unified model of cellular systems is a canonical challenge in biology. Recently, a wealth of public single-cell RNA sequencing data as well as rapid scaling of self-supervised learning methods have provided new avenues to address this longstanding challenge. However, rapid parameter scaling has been essential to the success of large language models in text and images, while similar scaling has not been attempted with Transformer architectures for cellular modeling. To produce accurate, transferable, and biologically meaningful representations of cellular systems, we develop AIDO.Cell, a pretrained module for representing gene expression and cellular systems in an AI-driven Digital Organism [1]. AIDO.Cell contains a series of 3M, 10M, 100M, and 650M parameter encoder-only dense Transformer models pre-trained on 50 million human cells from diverse tissues using a read-depth-aware masked gene expression pretraining objective. Unlike previous models, AIDO.Cell is capable of handling the entire human transcriptome as input without truncation or sampling tricks, thus learning accurate and general representations of the human cell’s entire transcriptional context. This pretraining with a longer context was enabled through FlashAttention-2, mixed precision, and large-scale distributed systems training. AIDO.Cell (100M) achieves state-of-the-art results in tasks such as zero-shot clustering, cell-type classification, and perturbation modeling. Our findings reveal interesting loss scaling behaviors as we increase AIDO.Cell’s parameters from 3M to 650M, providing insights for future directions in single-cell modeling. Models and code are available through ModelGenerator in https://github.com/genbio-ai/AIDO and on Hugging Face.

## 1 Introduction

Emergent behavior at scale is a common feature of biology. Cellular systems are often conceptualized as a series of chemical reactions, where many molecular interactions – carefully tuned over billions of years of evolution – give rise to robust, redundant, and responsive behaviors. From this visual, a unified model of cellular dynamics is both intuitive and plausible yet obviously intractable. Such a model is a grand challenge of modern biology [2, 3], which would revolutionize our ability to understand and manipulate cells, enabling drug discovery, personalized medicine, and fundamental knowledge about life.

Historically, the intractability of this goal required the use of statistical and mathematical methods with limited scope, adopting a one-model-one-task approach to research to characterize cellular systems piecewise. This is reminiscent of a bygone era of natural language processing, with bespoke models for sentiment analysis, embedding, grammar, etc. Now, large language models (LLMs) form a common foundation for all language tasks, in many cases drastically improving performance and transferability through the simple principles of scaling and unsupervised learning [4, 5, 6, 7].

In recent years, a series of breakthroughs in both high-throughput experimental biology and the development of large pretrained models on transcriptomes have fueled optimism on the possibility of similar models for cellular biology. High throughput biological screens have led to the collection of massive datasets spanning different cells and tissue systems. On the experimental side, there has been an exponential increase in rich and diverse reference datasets dedicated to comprehensively charting the landscape of various cellular systems[8, 9, 10], tissues, and perturbations [11, 12, 13]. On the modeling side, recent cellular FMs have demonstrated the capacity to cluster new cell types [14, 15], elucidate gene-gene interactions [16], and accurately predict combinations of gene knockouts by pretraining on millions of transcriptomes [17, 18].

Despite these successes, there still remains a significant gap between hand-crafted models and these transcriptome FMs. Linear models are often comparable to or outperform existing FMs [19, 20, 21, 22]. Towards this direction, we revisited reasons why existing models underperform. The first likely reason being scale with respect to both the model and data, as LLMs in NLP only outperformed supervised counterparts after scaling to billions of parameters.

Another possible reason is that the masked language modeling (MLM) objective in many models may be modeling an incomplete gene expression distribution due to down-sampling or truncation of genes to fit smaller context lengths. This truncation of the transcriptome can hinder the model’s ability to capture complex interactions from repressed genes and learn the true conditional gene expression distribution through MLM. Nonetheless, modeling the entire 20K-gene transcriptome is challenging because the Transformer’s attention mechanism has quadratic complexity with respect to context length. While scBERT and scFoundation use memory-efficient architectures to approximate attention, models like Performer [23] do not exhibit the same scaling laws as Transformers [24]. Moreover, subquadratic architectures may involve representational trade-offs [25] and exhibit explicit inductive biases [24].

In light of these challenges, we developed AIDO.Cell, a large-scale single-cell FM designed to prioritize learning rich representations for *human cells* at scale. Unlike previous models, AIDO.Cell is capable of handling the entire transcriptome (19,264 context length) as input without truncation or sampling tricks, thus learning accurate and general representations of the cell’s entire transcriptional context. We use the read-depth-aware (RDA) pretraining objective [17], which uses the masked gene expression of low read-depth to predict the expression of high read-depth. Our single-cell FM has been pre-trained at scale on over 50 million cells using the RDA pretraining objective. We employed large-scale distributed training strategies powered by state-of-the-art machine learning systems [26, 27] and FlashAttention2 [28] for high-throughput training across hundreds of GPUs, producing a series of single-cell FMs at 10M, 100M, and 650M parameter sizes. We found that training a model with a dense transformer backbone not only achieves state-of-the-art performance on various downstream tasks but also demonstrates strong scaling laws with interesting behavior as we scale the model’s parameters.

## 2 Methods

At its core, AIDO.Cell uses the bidirectional Transformer encoder-only architecture (BERT) [29], with several enhancements to improve its suitability for single-cell data. The read-depth-aware [17] pretraining objective encourages the model to learn a representation that is both robust to high variance in read-depth, and process gene expression directly as continuous values. Utilizing large-scale distributed pretraining [26] and systems-level optimizations such as FlashAttention-2 [28], we were able to train AIDO.Cell at scale.

**Figure 1:**
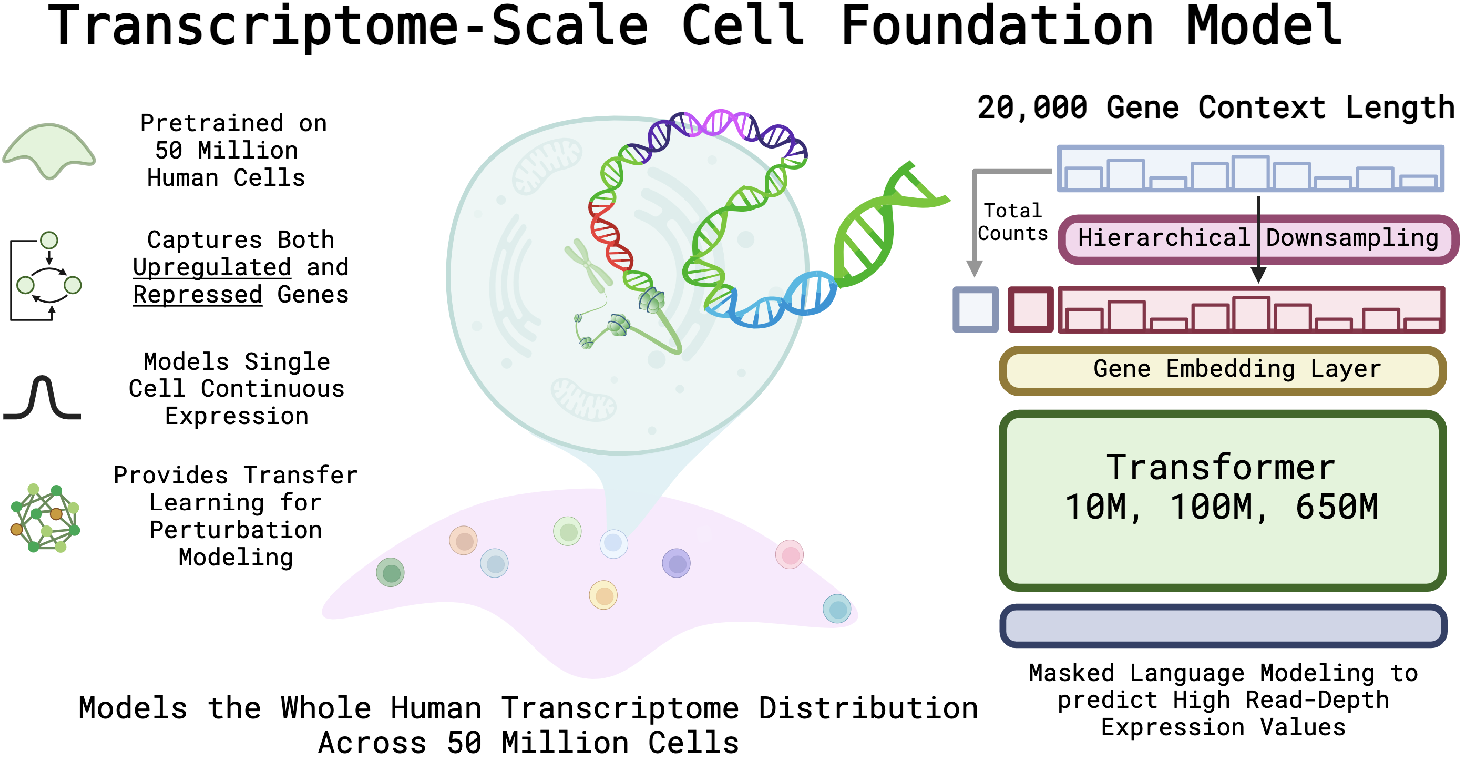
Overview of AIDO.Cell. Leveraging modern large-scale distributed training techniques and machine learning systems, we are able to scale the dense Transformer architecture up to 650M parameters over the human transcriptome of 20K genes.

**Table 1:** A list of a few representative single-cell foundation models and their properties.

| Name | Year | Architecture | Context | Whole Transcriptome | Params |
| --- | --- | --- | --- | --- | --- |
| scBERT[14] | 2022 | Performer | 16902 | Yes | 100M |
| GeneFormer[16] | 2023 | Transformer | 1024 | No | 10M |
| scGPT[18] | 2023 | Transformer | 1200 | No | 100M |
| scFoundation | 2023 | Trans. + Perf. | 2048/19264 | Yes | 120M |
| UCE[15] | 2024 | Transformer | 1024 | No | 650M |
| AIDO.Cell | Ours | Transformer | 19264 | Yes | 3/10/100/650M |

### 2.1 AIDO.Cell Architecture

#### 2.1.1 Gene Expression Embedding

AIDO.Cell uses the auto-discretization strategy from xTrimoGene [30] to effectively represent continuous values at high resolution. Previous works have shown that conventional ways of binning the gene expression input into discrete tokens, such as rounding to the nearest integer or calculating fixed bins based on frequency, have a detrimental effect on performance [30]. The auto-discretization strategy transformers each of the continuous expression values into a weighted linear combination of *b* learned tokens embeddings. The purpose of this is to allow the model to learn a flexible but shared embedding for each expression value.

First, the strategy initializes a random look-up table *T* ∈ R^*d×b*^, where *d* is the embedding dimension and *b* is the number of “tokens”. Then, we transform the expression value *v* ∈ R into a series of weights *α* ∈ R^*b*^ through a two-layer feed-forward (parameterized by **W**_1_, **W**_2_) and LeakyReLU in between, resulting in: (*α* = Softmax(**W**_2_ · LeakyReLU(**W**_1_ · *v*))). Then, the final output representation for each continuous value is a weighted linear combination of the “tokens” in the lookup table (*x* = *α* · *T*). We also learn dedicated embeddings for *mask token*, which we directly apply after the Gene Expression Embedding module.

#### 2.1.2 Transformer Backbone

AIDO.Cell uses a standard dense Transformer as its backbone [31]. The primary motivation for using a dense transformer as opposed to other memory-efficient architectures is to reduce the inductive bias and learn data-driven representations with minimal assumptions from the architecture. Many existing linear attention methods have are either designed for causal modeling [32] and/or have an inductive bias on a sequential ordering to the input [32, 33, 34]. Furthermore, existing studies have shown that many methods that approximate attention, such as Performer, do not necessarily exhibit the same scaling behaviors compared to attention [24]. With the goal of both scaling and minimizing inductive biases in the underlying data structure, we employed full dense attention. Pretraining at full dense attention was made possible through FlashAttention2 [28] and BF16 precision [35]. FlashAttention-2 employs tiling and block-wise computations to partition the QKV matrices into blocks that can be processed in SRAM (static random access memory). This, along with many other optimizations, allows for dense attention to be linear in memory instead of quadratic with respect to sequence length.

Although this does incur a significant computational cost for pretraining due to the quadratic complexity, ideally after this one-time cost, the model can be directly used or efficiently finetuned using methods such as LoRA [36] on various downstream tasks.

Our model architecture details can be found in 2. Our architectures employ LayerNorm [37] and SwiGLU [38], with no attention or MLP dropout [39], to increase model capacity.

**Table 2:**
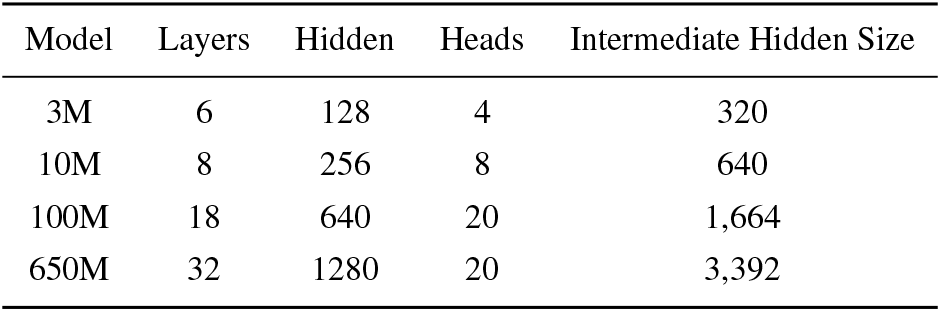
Model Architecture Parameters of AIDO.Cell.

| Model | Layers | Hidden | Heads | Intermediate Hidden Size |
| --- | --- | --- | --- | --- |
| 3M | 6 | 128 | 4 | 320 |
| 10M | 8 | 256 | 8 | 640 |
| 100M | 18 | 640 | 20 | 1,664 |
| 650M | 32 | 1280 | 20 | 3,392 |

### 2.2 Training Procedure

For training, our model underwent three epochs of the full pretraining dataset. We used a cosine learning rate scheduler with a linear warm-up of 5% for 150,000 iteration steps in total. We used a max learning rate of 3e-4 for the 100M and the 650M models. We used the AdamW optimizer with *β*_1_ = 0.9 and *β*_2_ = 0.95 and weight decay of 1e-2. We trained our models with bfloat-16 precision to optimize on memory and speed. The training took place over 256 H100 GPUs over three days for the 100M, and eight days for the 650M. Despite the long pretraining context-length, we did not need to employ tensor or pipeline parallelisms in these models.

We decided to train our model with a batch size of 1024 transcriptomes per step. Although traditionally, for masked-language-modeling, around 2 or 4 million tokens are used in a batch [40], because of the sparsity in single-cell data, we decided to make sure each batch had 1024 samples. Each sample has a full 19264 gene sequence length. Assuming a single-cell sparsity of around 10 ∼ 20%, this resulted in around 2 to 4 million nonzero genes processed per batch.

### Pre-training framework and parallelism

Our pre-training framework is built on the Megatron-LM Core version [41, 27], which is powered by the Transformer Engine [42]. Since pre-training of all AIDO.Cell models fits on a single GPU, we employ data parallelism across our 256-GPU cluster to ensure communication efficiency and good GPU utilization.

### 2.3 Pretraining Data

AIDO.Cell was pretrained on a diverse dataset of 50 million cells from over 100 tissue types. We followed the list of data curated by scFoundation in the supplementary[17]. This list includes datasets from the Gene Expression Omnibus (GEO) [9], the Deeply Integrated human Single-Cell Omnics data (DISCO)[43], the human ensemble cell atlas (hECA) [44], Single Cell Portal [45] and more. After preprocessing and quality control, the training dataset contained 50 million cells, or 963 total billion gene tokens. We partitioned the dataset to set aside 100,000 cells as our validation set.

#### 2.3.1 Read Depth Aware Pretraining Objective

As seen in existing works, there appears to be a trade-off between robustness to technical effects and representational accuracy. For example, the authors of GeneFormer have noted that although the normalized rank-value encoding allows for strong robustness to batch effects, the encoding does not fully take advantage of the precise gene measurements provided in transcript counts [16]. On the other side, the raw gene expression can have a high technical variation that can confound analyses, such as high variance in sequencing read depth, which is the difference in total read count between different experimental setups.

The read-depth-aware (RDA) pretraining objective [17] was first employed by scFoundation and aims to have the model predict the gene expression of high read depth from gene expression of low read count depth. To construct low read-depth samples, we first half the cells randomly during training, and then sample from a beta-binomial distribution, specifically:

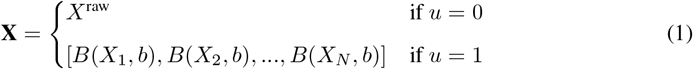

where *u* ∼ *Bernoulli*(0.5) and *b* ∼ *Beta*(2, 2). This was done to ensure that the expected log fold change between *X*_raw_ and the model input **X** is fixed to 1*/b*, where *b* ∼ *Beta*(2, 2). We then concatenate two total counts, *S* and *T* to the input. *S*, the “source” is the total read count (log normalized) of **X**. *T*, the “target”, is the total read count (log normalized) of *X*_raw_. Therefore, the input to the model is **X**^input^ = concat[**X**, *T, S*]. The goal was to train the model to predict the higher read count depth *X*_raw_ based on **X**_input_. Specifically, the mean squared error (MSE) loss on masked values 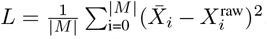, where 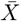 be the output vector of AIDO.Cell and *M* is the set of masked positions. This is the same pretraining objective used by scFoundation. In training, we mask out 30% of nonzero values, and 3% of zero values. Roughly speaking, there are 10 times more zeroed gene expression values than nonzero, so this masking ratio was to have roughly an equal proportion of masked nonzero and zero values.

## 3 Results

### 3.1 Scaling Results For Single Cell

The majority of existing studies on scaling laws are explicitly built around cross-entropy loss [46, 40], and it can be challenging to extrapolate training dynamics to different pretraining objectives. Masked MSE loss represents a substantial divergence from past work, but a necessary design choice for single cell gene expression. To determine compute-optimal scaling on single-cell data, a key milestone on the path to accurate and general single cell foundation models, we conduct a new scaling study on AIDO.Cell (Fig. 3.1). In this study, we trained AIDO.Cell at 3M, 10M, 100M, and 650M parameter scales to identify computational asymptotes. We adopted a similar procedure to that of [47], where we kept all hyperparameters the same and only scaled up the model size. In all cases, the model predicts masked expression values over the entire transcriptome.

**Figure 2:**
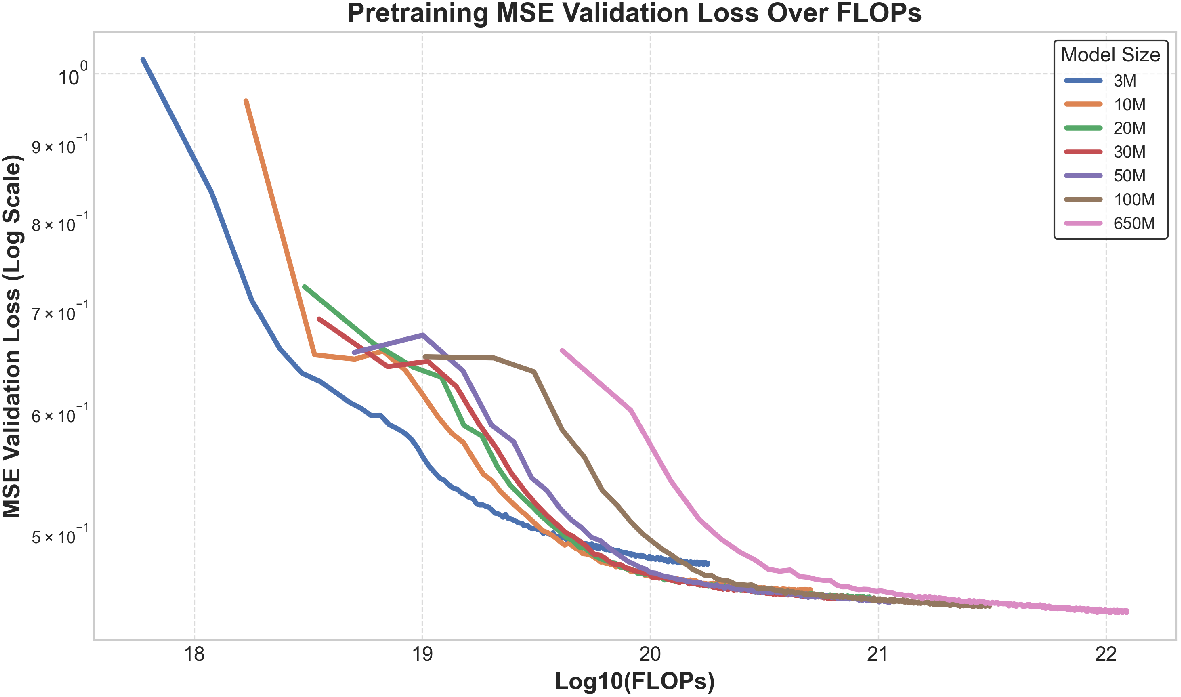
Pretraining Validation Loss Across 3M, 10M, 20M, 30M, 50M, 100M, and 650M

**Figure 3:**
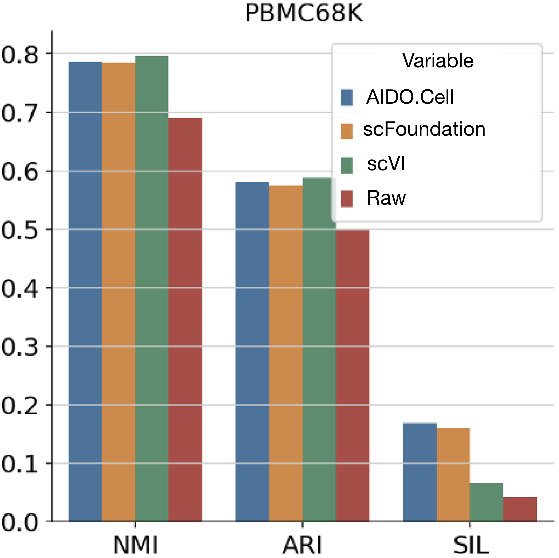
Gene expression foundation model embeddings evaluated on zero-shot cell-type-specificity clustering metrics.

We conducted experiments over model sizes of 3M, 10M, 20M, 30M, 50M, 100M, and 650M to observe of the scaling behavior of AIDO.Cell over the 50 Million single-cell dataset. Overall, we observed that scaling the model size improves both training and validation loss. However, in our dataset, we do not observe the same clear scaling power laws observed in NLP. It appears that after our 100M size, the improvement in validation loss is not as large as previous models before. There are a few possible reasons to this. The first being that this may result from a high irreducible loss due to biological or experimental noise. The second reason is that the objective of mean-squared error may scale in a less favorable manner compared to that of cross-entropy. Overall, this provides hints into how we should revisit data curation for single-cell foundation models, highlighting the need for more metrics to measure both dataset quality and diversity for pretraining.

The following downstream evaluations will primarily be focused on the AIDO.Cell (100M). This is to compare with the existing SOTA models at comparable parameter sizes. We plan to evaluate our large model AIDO.Cell (650M) in our next steps.

### 3.2 Zero-Shot Clustering Results

For zero-shot clustering, we leveraged the cell embeddings obtained from our 100M pre-trained checkpoint and compared them against previous baselines: scFoundation [17] and scVI [48] on a PBMC dataset from Zheng68K [49]. Using these embeddings, we followed scFoundation [17] and applied the Leiden clustering algorithm to identify distinct cell populations without any fine-tuning. We evaluated the performance of the clustering using multiple metrics, including Adjusted Rand Index (ARI) and Normalized Mutual Information (NMI), which assess the agreement between the predicted clusters and ground truth labels. Additionally, Silhouette Score (SIL) was used to measure the internal consistency of the clusters.

Figure 3 shows the zero-shot clustering results for our model, and Figure 4 shows a qualitative UMAP comparison. In the UMAP comparisons, AIDO.Cell (b) reveals clear, well-separated clusters representing various immune cell types, including B cells, T cell subtypes, and monocytes. These clusters align well with known biological labels, outperforming scFoundation, which exhibits more overlap between cell types. Quantitatively, our approach achieves higher scores across all metrics, as shown in Figure 3. The improved NMI and ARI reflect the model’s superior alignment with true cell types, while the higher SIL indicates better internal cluster consistency. These results suggest that our dense representations capture biologically meaningful structures at the single-cell level, even without task-specific fine-tuning, showing the effectiveness of our method for zero-shot clustering.

**Figure 4:**
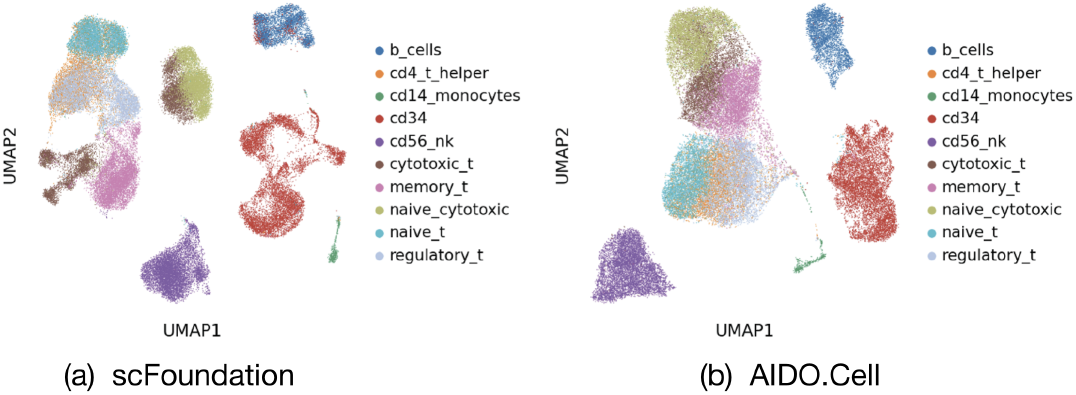
UMAP visualization of gene expression foundation model embeddings, colored by cell type.

### 3.3 Cell Type Classification Task Results

Cell type annotation has been one of the most classic and essential tasks in the field of single-cell research. To evaluate AIDO.Cell in terms of the quality of single-cell encoding, we carried out experiments by finetuning and evaluating the model on two datasets: 1. Zheng68K [49], a classic PBMC dataset widely used for the cell annotation task benchmarking; 2. Segerstolpe [50], a small pancreas dataset that have shown to be challenging as for cell annotation. We used AIDO.Cell (100M) for a close parameter comparison.

To make a fair comparison to scFoundation and other baseline models, we followed the same architecture as scFoundation for downstream classification by adding a two-layer MLP head on top of the encoder of AIDO.Cell. This MLP head is fine-tuned together with the last encoder layer. We also used the same weighted cross entropy loss as scFoundation to mitigate the impact of imbalanced target classes:

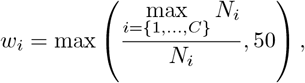

where *w*_*i*_ is the corresponding weight for cell type *i* and *N*_*i*_ is the number of cells of cell type *i* in the training set. In addition, to maintain consistency with the reported results of scFoundation, we used the same data splits as well as macro F1 as the evaluation metric.

Our final results on the test set is reported in table 3. We have performed a small-scale hyperparameter sweep by training with the combination between learning rate of 1e-3 and 1e-4 as well as effective batch size of 128 and 256. For Segerstolpe, due to the small training set and consequently higher risk of overfitting, we performed additional sweep experiments with regularization-related hyperparameters, including dropout rate of 0.5 and weight decay of 1e-2. The final model, which is chosen based on the macro F1 score on the validation set have outperformed other models on Zheng68K and achieved comparable results to scFoundation on Segerstolpe.

**Table 3:** Comparison of model performance on Zheng68K and Segerstolpe datasets.

| Model | Zheng68K<br>F1 Macro | Segerstolpe<br>F1 Macro |
| --- | --- | --- |
| AIDO.Cell (100M) | **0.761** | 0.910 |
| scFoundation (100M) | 0.736 | **0.914** |
| scBERT | 0.67 | 0.67 |
| CellTypist | 0.725 | 0.812 |
| scANVI | 0.395 | 0.521 |
| ACTINN | 0.649 | 0.722 |
| Scanpy | 0.547 | 0.54 |
| SingleCellNet | 0.598 | 0.806 |

### 3.4 Perturbation Modeling Task Results

One of the goals of cellular modeling is to understand, reason, and ultimately design interventions to reprogram cellular behavior, paving the path toward personalized medicine. This has become more feasible with the strides made in both single-cell sequencing and CRISPR gene editing technologies, which allows for a precise understanding of the causal effects when applying a treatment to a control population of cell lines. However, the vast array of possible perturbation combinations is countably too large to perform experimentally but can be traversed computationally. GEARs is a graph convolutional network constructed from gene-ontology and gene co-expression networks to predict the change in expression from single or combinations of perturbations.

Following the task formulation in both GEARs[51] and scFoundation[17], we combine GEARs and AIDO.Cell (100M) to predict genetic perturbations. One of the primary strengths of cellular foundation models is their ability to contextualize cells with their rich internal representation. These contextualized representations can improve the downstream performance of existing models for challenging tasks such as cellular perturbation prediction.

AIDO.Cell takes as input a cell and provides a latent representation for each gene in the cell. This representation is used to parameterize the nodes of GEARs and make predictions on changes in gene expression from single or combinations of genetic perturbations. Our results for the Norman et al. dataset are in table 4).

**Table 4:**
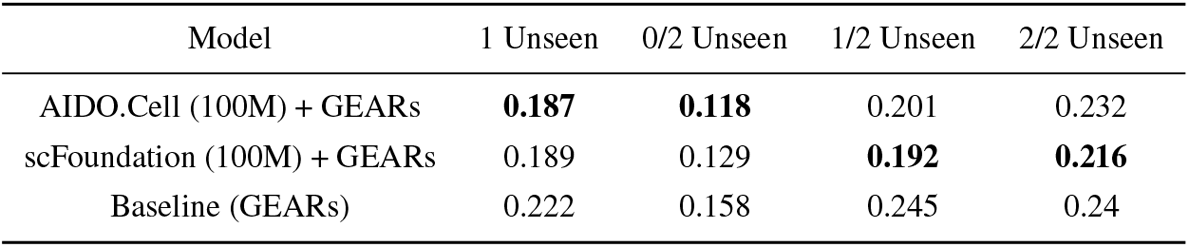
MSE on Differentially Expressed Genes in the Norman et al. Dataset.

| Model | 1 Unseen | 0/2 Unseen | 1/2 Unseen | 2/2 Unseen |
| --- | --- | --- | --- | --- |
| AIDO.Cell (100M) + GEARs | **0.187** | **0.118** | 0.201 | 0.232 |
| scFoundation (100M) + GEARs | 0.189 | 0.129 | **0.192** | **0.216** |
| Baseline (GEARs) | 0.222 | 0.158 | 0.245 | 0.24 |

We followed the same perturbation formulation to scFoundation. For scFoundation+GEARs and default baseline GEARs, we used the same hyperparameters referenced in the paper to reproduce their results. For AIDO.Cell, We performed a small hyper-parameter sweep over the learning rate and batch size, where we varied the batch size to be 30 or 60 and the learning rate to be either 1e-2 or 1e-3. This was based on the hyperparameters used by scFoundation, which are a batch size of 30 and a learning rate of 1e-2. Following the same framework, we picked the model with the best validation score over differentially expressed genes and reported the same metrics to scFoundation. We found that AIDO.Cell reaches SOTA performance, outperforming GEARs, and achieves comparable scores to scFoundation over a short hyperparameter search.

## 4 Discussion

Large-scale deep learning, pioneered in natural language processing, has shown surprising and emergent behaviors which have been difficult to replicate in other domains. While it is often viewed as safest to retain the design choices of previous works when translating to a new domain, the underlying model and the architectural alignment with the physical mechanisms of the data have to be the primary consideration. Simply put, cells are not sentences. Using the same architectures, objectives, and context lengths as LLMs has not led to accurate and general models of cellular biology [20]. Nonetheless, foundation models must scale favorably with respect to data, size, and compute [46]. Many single-cell foundation models have been proposed [18, 52, 16, 14, 53, 54], but rigorous studies on the effect of scaling laws, architectures, and pretraining objectives remain sparse or closed-source [55]. To address this, we propose AIDO.Cell, a scalable transformer-based foundation model for single cell gene expression, and a component of a larger AI-driven Digital Organism [1]. Leveraging both highly scalable architectures and biologically motivated design choices that deviate from previous works on LLMs, AIDO.Cell achieves state-of-the-art performance on canonical benchmarks in transcriptomics. Overall, AIDO.Cell promises a platform which benefits from innovations in high throughput data collection in biology and large-scale distributed training in natural language processing to explore the scaling properties of pretraining and finetuning at unprecedented scale and speed.

